# Differentiating amyloid beta spread in autosomal dominant and sporadic Alzheimer’s disease

**DOI:** 10.1101/2021.06.25.449939

**Authors:** Elizabeth Levitis, Jacob W. Vogel, Thomas Funck, Vladimir Halchinski, Serge Gauthier, Jonathan Voglein, Johannes Levin, Tammie Benzinger, Yasser Iturria-Medina, Alan C. Evans, for the Dominantly Inherited Alzheimer Network, for the Alzheimer’s Disease Neuroimaging Initiative

## Abstract

Amyloid-beta (Aβ) deposition is one of the hallmark pathologies in both sporadic Alzheimer’s disease (sAD) and autosomal dominant Alzheimer’s disease (ADAD), the latter of which is caused by mutations in genes involved in Aβ processing. Despite Aβ deposition being a centerpiece to both sAD and ADAD, some differences between these AD subtypes have been observed with respect to the spatial pattern of Aβ. Previous work has shown that the spatial pattern of Aβ in individuals spanning the sAD spectrum can be reproduced with high accuracy using an epidemic spreading model (ESM), which simulates the diffusion of Aβ across neuronal connections and is constrained by individual rates of Aβ production and clearance. However, it has not been investigated whether Aβ deposition in the rarer ADAD can be modeled in the same way, and if so, how congruent the spreading patterns of Aβ across sAD and ADAD are. We leverage the ESM as a data-driven approach to probe individual-level variation in the spreading patterns of Aβ across three different large-scale imaging datasets (2 SAD, 1 ADAD). We applied the ESM separately to the Alzheimer’s Disease Neuroimaging initiative (N=737), the Open Access Series of Imaging Studies (N=510), and the Dominantly Inherited Alzheimer’s Network (N=249), the latter two of which were processed using an identical pipeline. We assessed inter- and intra-individual model performance in each dataset separately, and further identified the most likely epicenter of Aβ spread for each individual. Using epicenters defined in previous work in sAD, the ESM provided moderate prediction of the regional pattern of Aβ deposition across all three datasets. We further find that, while the most likely epicenter for most Aβ-positive subjects overlaps with the default mode network, 13% of ADAD individuals were best characterized by a striatal origin of Aβ spread. These subjects were also distinguished by being younger than ADAD subjects with a DMN Aβ origin, despite having a similar estimated age of symptom onset. Together, our results suggest that most ADAD patients express Aβ spreading patters similar to those of sAD, but that there may be a subset of ADAD patients with a separate, striatal phenotype.

## Introduction

To date, there is no cure for Alzheimer’s disease (AD), the principle neurodegenerative cause of dementia. Treating patients with dementia is costly - in 2009, the average cost for a patient with AD was roughly 57,000 USD^1^. The socioeconomic gravity of treating AD has spurred research seeking to prevent or mitigate AD by first developing biomarkers that can be used for early diagnosis and monitoring^2^. The two main pathological signs of AD are neurofibrillary tau tangles and Aβ senile plaques, and both are required to definitively confirm AD at autopsy^3^. Most hypothetical models of AD progression have been rooted in the amyloid cascade hypothesis, which posits that excessive amounts of soluble Aβ cause a buildup of insoluble Aβ, disrupting synaptic function and accelerating tau hyperphosphorylation^4^. Most cases of AD are sporadic in nature, and the much rarer autosomal dominant form of AD is caused by mutations in genes - namely, APP, PSEN1, PSEN2 - that impact the processing of the amyloid precursor protein from which the Aβ peptide is cleaved. While ample research has pointed to accumulation of Aβ in the brain as being one of the earliest pathological biomarkers in both sAD and ADAD, we know quite little about where and how Aβ begins to accumulate, how it spreads in the brain, and whether either of these is variable across individuals. An ADAD mutation virtual guarantees amyloidosis, making carriers of these mutations incredibly important for the study of amyloid-related processes and brain changes in AD. However, it is still unclear just how similar ADAD and sAD are with respect to the progression of various biomarkers, including Aβ.

In general, most studies in this domain have focused less on inter-individual variability and have primarily reported group differences. Unlike in sAD, where Aβ deposition is highest in neocortical areas, several groups have reported significantly increased striatal, thalamic, and neocortical Aβ deposition in ADAD mutation carriers compared with noncarriers^5,6^. One study evaluating differences between the PSEN1, PSEN2, and APP ADAD mutation types found that all mutation types had high striatal PiB binding while some mutation carriers had higher cortical PiB binding. Interestingly, PiB binding in the cortex was found to be lower in ADAD mutation carriers than age-matched subjects with probable sAD^7^. While the sample size of this study was small (n=30 ADAD mutation carriers, n=30 sAD subjects), the findings suggest that the most probable area(s) of earliest Aβ accumulation may not be homogenous amongst all ADAD mutation carriers.

Recently, an event-based model of disease progression was applied to ADAD mutation carriers. The authors found that the biomarker likeliest to exhibit the earliest deviation from normal levels was a cortical Aβ deposition measure, followed by Aβ deposition in the caudate, putamen, accumbens, and thalamus^8^. In sAD, a separate model leveraged CSF and Aβ signals to stage subjects according to Aβ accumulation status. In this study, subjects who were both CSF and Aβ negative according to a set of data-driven thresholds were deemed to be non-accumulators, whereas those who were CSF positive but Aβ negative were deemed to be early Aβ accumulators^9^. Regions pinpointed as areas of earliest accumulation were those that had significantly increased Aβ signal in early accumulators compared with non-accumulators. According to this system, the precuneus, medial orbitofrontal cortex, and posterior cingulate were all categorized as regions of early accumulation whereas the caudal anterior cingulate was pinpointed as an area of intermediate accumulation. Together, these studies suggest possible differences between ADAD and sAD in the earliest regions to accumulate Aβ.

While both data-driven approaches can be used to glean the order in which biomarkers can be detected at either a regional or global level, neither of them is mechanistic in nature. To better understand how Aβ or tau spreads in the brain, we can instead turn to an epidemic spreading model (ESM) developed to stochastically reproduce the propagation and deposition of misfolded proteins such as Aβ, tau, and alpha-synuclein. The overarching nonlinear differential equation of the model posits that the change in misfolded protein deposition in each macroscopic region of interest (ROI) is equal to the probability of endogenously producing and exogenously receiving misfolded proteins from connected ROIs, minus the probability of clearing the deposited misfolded proteins. This approach has previously been applied to model the spread of Aβ and tau across anatomical connections in individuals along the sAD spectrum^10,11^. When applied to over 700 subjects in the ADNI dataset, the ESM was able to explain 46-57% of the variance in the mean regional Aβ deposition probabilities of the distinct clinical subgroups and identified the posterior and anterior cingulate cortices as the seed regions of Aβ propagation. These seed regions are in agreement with what has been established in the literature^12^. Using the ESM, we can evaluate whether there is sufficient evidence to suggest that Aβ spreads along neuronal connections in ADAD as well. Furthermore, we can evaluate how similar ADAD and sAD are with respect to which regions Aβ begins spreading from. We tackle this question by applying the ESM within three different datasets representing sAD and ADAD, to both evaluate differences between ADAD and LOAD, as well as validate the previously published results in an independent dataset.

## Methods

### Participants

Participants for this study are comprised of individuals from three multi-center studies: the Dominantly Inherited Alzheimer’s Network (DIAN; https://dian.wustl.edu), the Alzheimer’s Disease Neuroimaging Initiative (ADNI; http://adni.loni.usc.edu), and the Open Access Series of Imaging Studies (OASIS; www.oasis-brains.org). While the ESM had already been applied to ADNI, a dataset representative of LOAD, we include an additional dataset for two reasons - (1) to validate the previously published results in an independent cohort and (2) to compare results in DIAN with a dataset that used the same radiotracer and was processed using the same pipeline.

The DIAN dataset represents individuals from families known to have mutations in the amyloid precursor protein (APP), presenilin 1 (PSEN1), and presenilin 2 (PSEN2) genes. Both mutation carriers and non-carriers were used for different stages of the analysis. We selected individuals who had at least one PIB PET scan and accompanying T1w scan from the 12th semiannual DIAN data freeze. For this study, DIAN serves as the dataset representative of ADAD.

The OASIS dataset is a compilation of participants from multiple studies, and the participants range from older, cognitively normal adults to those at various stages of cognitive decline and dementia.

### MRI and PET acquisition and preprocessing

MRI and PET acquisition procedures for the DIAN^13^, ADNI (http://adni.loni.usc.edu/methods/), and OASIS^14^ datasets have previously been described in detail.

It is important to note that the processing pipeline and the radiotracer for the ADNI dataset diverge from those used for DIAN and OASIS. For ADNI, the preprocessing pipeline is taken from the original ESM publication^10^. Briefly, individual AV45 PET scans were acquired and processed in the following order - dynamic co-registration, averaging across time, re-sampling and reorientation from native space to a standard voxel space, spatial filtering, and finally spatial normalization to MNI space. For the DIAN and OASIS dataset, whole-brain T1w scans and individual PiB-PET scans were acquired. Quality control was performed as per the ADNI protocol. FreeSurfer version 5.3 (http://surfer.nmr.mgh.harvard.edu) was used to derive subject-specific segmentations corresponding to regions in the Desikan-Killiany-Tourville atlas (DKT,^15^). Only cortical and subcortical regions overlapping with the Mindboggle DKT atlas were used, for a total of 78 regions^16^.

For both OASIS and DIAN, the PET Unified Pipeline (PUP; https://github.com/ysu001/PUP) was used to preprocess the PET scans. The processing steps used include smoothing, interframe motion correction and co-registration. Specifically, PET images in the 4dfp format are smoothed to achieve a common spatial resolution of 8mm to minimize inter-scanner differences (Joshi et al., 2009). PET-MR registration was performed using a vector-gradient algorithm (VGM) (Rowland et al., 2005). Co-registered summed PET scans in the 4dfp file format were downloaded from the CNDA portal (https://cnda.wustl.edu), and 4dfp images were subsequently converted to the Nifti file format for further analysis.

### Regional Aβ Probabilities

Traditionally, static PET processing involves quantifying co-registered PET images using standardized uptake value ratios (SUVR) for each ROI with respect to the average signal in a reference region devoid of specific tracer binding. The reference region typically used in AD Aβ PET imaging studies is the cerebellar cortex; however, amyloid deposition has been observed in the cerebellar cortex of individuals with ADAD^17^. Based on recent work seeking to clarify the optimal reference region for Aβ measurement using PiB-PET and the DIAN cohort, we used the brainstem as the reference region for the DIAN and OASIS datasets^18,19^. For the ADNI dataset, we used the Aβ deposition probabilities that had previously been generated (using a cerebellar reference region)^10^.

The original ESM paper introduced a voxelwise probability metric, which we will refer to as V ECDF, RR EVD. For each subject this approach creates a bootstrapped sampling consisting of 40,000 subsamples in the 5-95% of values in the reference region. Subsequently, an extreme value distribution (EVD) is created using the maximum value observed in each bootstrapped sample. The EVD is used to create an extreme cumulative distribution function, and for each voxel in the PET image, the probability of it being greater than every value in the EVD is computed. A final regional Aβ deposition probability is calculated as the average of the probabilities corresponding to each voxel within a given ROI. Given the overall higher PiB-PET signal in the brainstem than the cerebellar cortex, we use the 75th percentile value rather than the maximum in each bootstrap sampling to create the EVD when using the brainstem as the reference region.

For the DIAN dataset, we observed that noncarriers’ Aβ deposition probabilities were negligible in all ROIs except for the globus pallidus and thalamus, ROIs that have previously been observed to have nonspecific uptake of PiB^20,21^. Given their young age (Table 1), we are confident that the DIAN non carriers are truly amyloid-negative and are therefore a fully reliable control group. Subsequently, for each ROI, across all available timepoints, the noncarriers’ signal was used to create a ROI-specific control distribution. For each mutation carrier, we calculated a z-score for their Aβ binding probability in the ROI with respect to the ROI-specific control distribution. Within each ROI, we min-max scaled the absolute values of the z-scored signal across all timepoints to have probabilities in the [0,1] range again.

**Table 1:** Demographic information

| Dataset | DIAN |  | ADNI | OASIS |
| --- | --- | --- | --- | --- |
|  | T1 | T2 |  |  |
| N | 249 | 124 | 737 | 510 |
| Age (SD) | 39.01 (10.7) | 42.12 (9.7) | 72.43 (7.2) | 67.65 (9.8) |
| % Women | 56.3% | 60.1% | 44.9% | 57.8% |
| EYO (SD) | -8.54 (10.9) | -4.7 (9.8) | - | - |
| % ApoE4 | 30.1% | 29.53% | 51.7% | - |
| % Aβ Positive | 55% | 63.7% | 54% | 25% |
| % Cognitively Normal | 68.7% | 58.8% | 26.2% | 86.5% |

EYO = Estimated years to symptom onset; SD = Standard Deviation; T1 = Timepoint 1; T2 = Timepoint 2

### Epidemic Spreading Model

The spread of Aβ was simulated using the ESM, a diffusion model that has previously been used to simulate the spread of Aβ and tau in the ADNI dataset from an initial epicenter(s) and through an ROI network^10,11^. In addition to the connectivity between ROIs, subject specific propagation parameters influence the magnitude or extent of the spreading pattern. These parameters correspond to a global clearance rate, global production rate, and age of onset. These are fit by solving a non-linear differential equation designed to reproduce the overall regional pattern of Aβ deposition. The ESM is fit by searching the parameter space, and the set of parameters that yield the regional pattern of Aβ deposition most like the reference (observed) pattern is selected.

The main data input to the ESM is the ROI by Subject matrix reflecting regional Aβ deposition probabilities for each subject. Epicenters can either be supplied by the user or selected in a data-driven way. In the data-driven context, the ROI or combination of ROIs that best explain the average group-level pattern are returned as the epicenters. For a more detailed overview of the equations underlying the ESM, please refer to^10^.

### Connectivity Measures

In order to propagate Aβ signal across the brain, the ESM requires a matrix of pairwise relationships between ROIs. This informs the final regional pattern of Aβ. Earlier applications of the ESM tested whether Aβ spreads along synapses by using a structural connectivity matrix.

We used a structural connectivity matrix derived from diffusion spectrum imaging (DSI) scans of 60 young healthy subjects from the CMU-60 DSI template^22^. The acquisition and pre-processing steps have been described in detail in the original ESM paper and were based on methodology developed in an earlier paper^23,10^. Briefly, all images were non-linearly co-registered to MNI space, and orientation distribution functions (ODF) representing nerve fiber orientations were calculated. All intravoxel fiber ODF maps were averaged to create an ODF template, and an automated fiber tractography method was used to calculate probabilistic axonal connectivity values for each voxel and the surface of each grey matter region in the DKT atlas. Previously described anatomical connection probabilities were then generated for each ROI-ROI pair.

### Aβ Positivity

The ESM has previously been shown to be sensitive to spurious levels of signal, so we opted to confine our analysis to Aβ positive subjects^11^. We used gaussian mixture modelling to compute Aβ positivity thresholds in a data-driven way for both the PUP generated SUVRs and the probability values averaged across a composite set of regions that are implicated in AD. Specifically, these include the bilateral precuneus, superior frontal, rostral middle frontal, lateral orbitofrontal, medial orbitofrontal, superior temporal, and middle temporal ROIs. For each metric, we fit a two-component mixture model across the entire DIAN dataset - including non-carriers and mutation carriers - and estimated a cut-off. Only subjects who were positive on both the SUVR and probability metrics were considered Aβ positive for subsequent analyses. Since the DIAN and OASIS datasets were both processed using the WUSTL PET Unified Pipeline, we applied the cut-offs generated using the DIAN dataset to the OASIS dataset as well. Aβ positive subjects in these datasets were defined as those whose average Aβ value across a set of previously defined cortical areas surpassed 0.81 and 0.01236 for SUVR and deposition probability values, respectively. We illustrate the correspondence between within-subject composite Aβ SUVRs and deposition probabilities, as well as the GMM results in Figure S1. Aβ positive ADNI subjects were identified using a previously defined composite Aβ SUVR threshold of 1.11.

### Statistical analysis

Using the structural connectivity matrix and the cross-sectional baseline subject by region Aβ probability deposition matrix, we fit the ESM across different possible epicenters for the DIAN, ADNI, and OASIS datasets. Model performance for each experiment was evaluated by mean within-subject and global fit. Within-subject performance is evaluated as the Pearson r^2^ between the subject-specific observed and predicted regional Aβ deposition probabilities measured using PiB-PET. We evaluate global fit by averaging the observed and predicted regional Aβ probabilities across all subjects, respectively, and calculating the Pearson r^2^ between the averaged observed and predicted patterns. To ensure that our results are statistically significant and specific to the connectivity matrix we used, we scrambled the original connectivity matrix 100 times while preserving degree and strength distributions using the Brain Connectivity Toolbox (https://sites.google.com/site/bctnet/). We used the null distribution of the mean within-subject fit and global fit to calculate the mean and 95% confidence intervals for each ESM experiment.

Building off previous results suggesting that Aβ first accumulates in the posterior cingulate (PC) and caudal anterior cingulate (cAC) and subsequently spreads to other regions in the brain (in ADNI), we sought to evaluate whether this replicates in another LOAD dataset, as well as the DIAN dataset. Given our objective of evaluating whether a cortical or striatal epicenter better explains Aβ spreading patterns in ADAD, we repeated this analysis for all three datasets using the caudate and putamen as the seed regions. For a more data-driven approach to epicenter selection, in each dataset, we evaluated global fit using each bilateral ROI as an independent epicenter. For each subject we noted the epicenter that provided the best within-subject fit, and we assessed how frequently specific epicenters were present within each dataset. Given the lack of consensus about whether ADAD mutation carriers first accumulate Aβ in the striatum or neocortical regions that overlap with the default mode network, we further divided the possible epicenters into three subgroups - default mode network (DMN), striatum, and other. ROIs falling into the DMN group included the posterior cingulate, caudal anterior cingulate, rostral anterior cingulate, precuneus, and medial orbitofrontal cortex. The striatum subgroup included the caudate and putamen, and the other group contained all ROIs not in the other three subgroups. Using these data-driven epicenter subgroups, we compared within-subject model performance using either the caudate and putamen or cAC + PC as epicenters across the epicenter subgroups. We evaluated the statistical difference in the models’ performance across epicenter subgroups using the non-parametric Kolmogorov-Smirnov (K-S) test statistic.

After stratifying subjects across epicenter subgroups (DMN, Striatum, and Other), we examined associations with age and EYO. We additionally ran ordinary least-squares general linear models (GLMs) to assess the relationship between the epicenter subgroup and the Aβ signal in all ROIs while covarying for age and sex. We FDR corrected the relationships used the Benjamini–Hochberg approach.

As a follow-up, we evaluated the test-retest reliability of the best within-subject epicenter for each subject that had two scans.

## Results

### Sample information

Baseline PiB-PET scans measuring fibrillar Aβ load were available for 249 ADAD mutation carriers in the DIAN dataset. 124 of these mutation carriers had one follow-up PiB-PET scan, and 44 of them had two follow-up scans. Baseline AV45-PET scans were available for 737 individuals from the ADNI dataset, and baseline PiB-PET scans were available for 510 individuals from the OASIS cohort. Demographic information for this sample can be found in Table 1.

### Putative areas of early Aβ accumulation in LOAD do not explain the full picture in ADAD

To evaluate whether neuronal connectivity can explain the whole-brain pattern of Aβ in both ADAD mutation carriers and individuals from the OASIS dataset, we fit the ESM to regional Aβ deposition probabilities derived using PiB-PET or AV45-PET data (see Methods).

We first evaluated how well previously identified regions of early amyloid, namely cingulate and striatal regions, recapitulate group-level whole-brain Aβ patterns across all three datasets. We will refer to the model using the caudal anterior cingulate (CAC) and posterior cingulate (PC) as epicenters as the CAC + PC model, and the one using the caudate and putamen as the striatal model. In the DIAN dataset, the model using the CAC and PC as seed regions explained 27% (null model mean r2 [95% CI] = 0.119 [0.089, 0.164]; p < 0.01) of the aggregated pattern of Aβ, and on average explained 14.6% (null model mean r2 [95% CI] = 0.07 [0.002, 0.179]; p = 0.1) of the regional pattern of Aβ within individual subjects (Fig 1a). In Aβ positive subjects, the global fit and the mean within subject fit improved to 31% and 20.7% (p < 0.05), respectively. When stratifying performance across the three main mutation types, we found that there was no significant difference between the three groups. In line with the results that had been previously shown for the ADNI dataset in^10^, the CAC + PC model explained 53.9% (null model mean r2 [95% CI] = 0.103 [0.074, 0.148]; p < 0.01) of the aggregated pattern of Aβ and on average explained 39.1% (null model mean r2 [95% CI] = 0.087 [0.002, 0.217]; p < 0.01) of the regional pattern of Aβ in individual subjects. In Aβ positive subjects, the global fit and the mean within subject fit changed slightly to 51% and 38%, respectively.

**Figure 1.**
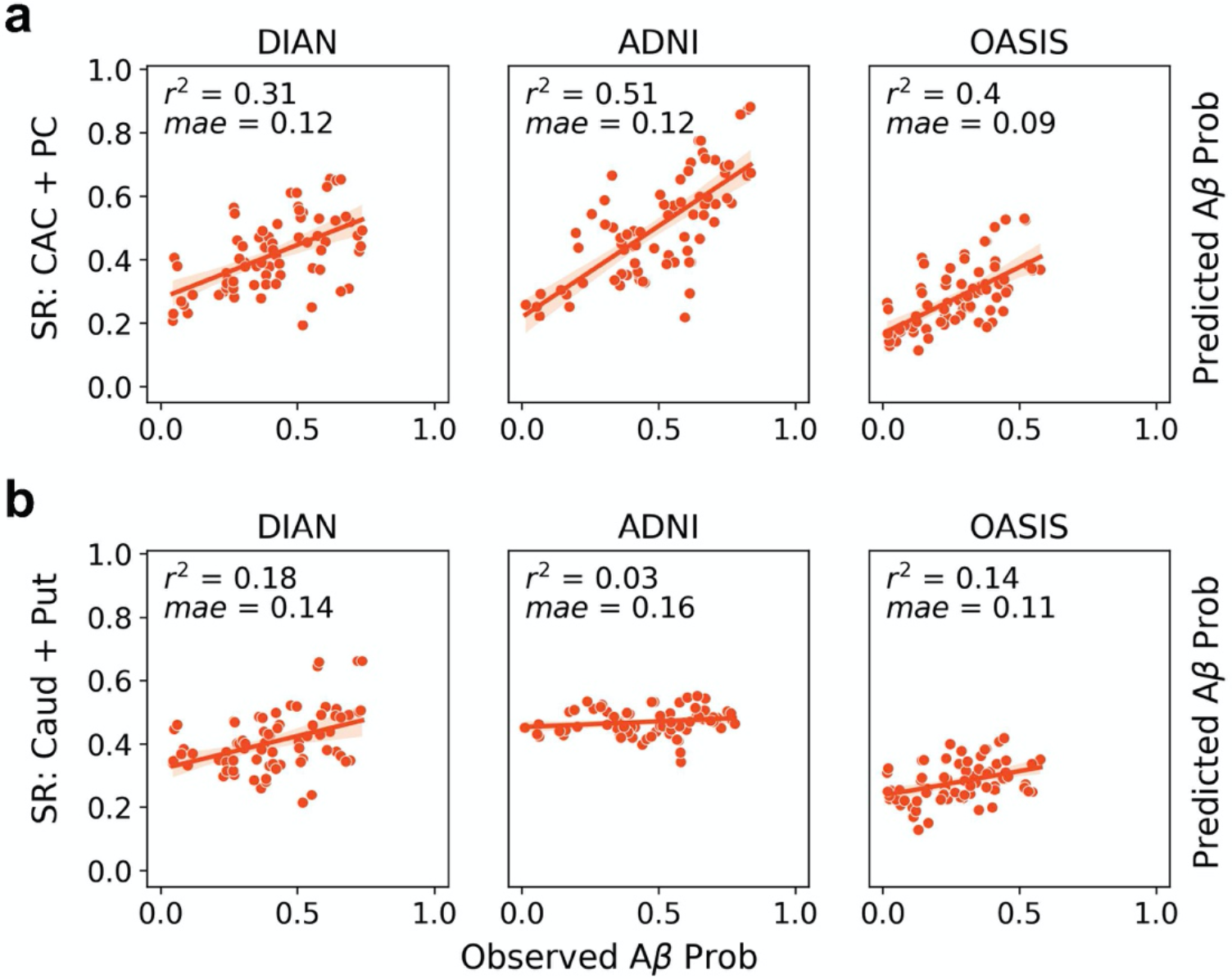
Comparison of global model fit across datasets and epicenters. ESM performance (global fit) across the ADNI, OASIS, and DIAN datasets using either the (a) posterior cingulate and caudal anterior cingulate or (b) caudate and putamen as epicenters. Each dot represents the observed and predicted mean signal for an ROI across all subjects within a dataset. Only Aβ positive subjects were included.

In the LOAD validation dataset, OASIS, the performance was lower than what had previously been reported for ADNI. Across the whole dataset, the CAC + PC model explained 28% (null model mean r2 [95% CI] = 0.158 [0.123, 0.217]; p < 0.01) of the aggregated pattern of Aβ and on average explained 9% (null model mean r2 [95% CI] = 0.063 [0.017, 0.139]; p = 0.15) of the within subject variance. However, when we only look at Aβ positive individuals, the global fit and the average within subject fit increased to 40% (null model mean r2 [95% CI] = 0.14 [0.098,0.18]; p < 0.01) and 21% (null model mean r2 [95% CI] = 0.082 [0.002, 0.196]; p = 0.04), respectively, and the results were significant.

Since a primary goal of this study was to identify whether a cortical or striatal epicenter better explains the regional patterns of Aβ in DIAN, we additionally repeated the same analysis using the caudate and putamen as the seed regions. When applied to ADNI, the striatal model performed poorly. It explained 3% (null model mean r2 [95% CI] = 0.055 [0.043, 0.072]; p = 1) of the aggregated pattern of Aβ and on average explained 5% (null model mean r2 [95% CI] = 0.05 [0.001, 0.139]) of the within-subject Aβ patterns in Aβ positive subjects. In DIAN Aβ positive subjects, the striatal model explained 18% (null model mean r2 [95% CI] = 0.103 [0.072, 0.146]; p < 0.02) of the aggregated pattern of Aβ and on average explained 17.2% (null model mean r2 [95% CI] = 0.085 [0.001, 0.256]) of the within-subject pattern. In Aβ+ OASIS subjects, the striatal model explained 14% (null model mean r2 [95% CI] = 0.084 [0.062, 0.133]; p < 0.02) and on average 11.4% (null model mean r2 [95% CI] = 0.059 [0.001, 0.16]; p = 0.15) of the global and within-subject results, respectively.

### Epicenter heterogeneity in DIAN compared with ADNI and OASIS

We initially compared ESM performance between the ADNI and DIAN dataset using a priori defined epicenters. Next, we ran the ESM using each bilateral ROI as the model epicenter to evaluate which ROI best explains the whole-brain patterns of Aβ in each dataset. We assigned each participant to an epicenter subgroup based on which ROI yielded the best within-subject performance (as described in 2.7).

In Figure 2a-b we show the relative breakdown of epicenter subgroups within the datasets in all subjects, and in Aβ positive subjects only. In Aβ positive subjects from ADNI and OASIS, most subjects have an epicenter in the DMN, while the remaining subjects fall into the Other category. Specifically, 89.2% of Aβ positive ADNI subjects and 72.7% of Aβ positive OASIS subjects have a DMN epicenter. In the DIAN dataset, there was substantially more heterogeneity, with 59.1% of Aβ subjects falling into the DMN group, 13.1% into the striatum group, and 27.7% into the Other group.

**Figure 2.**
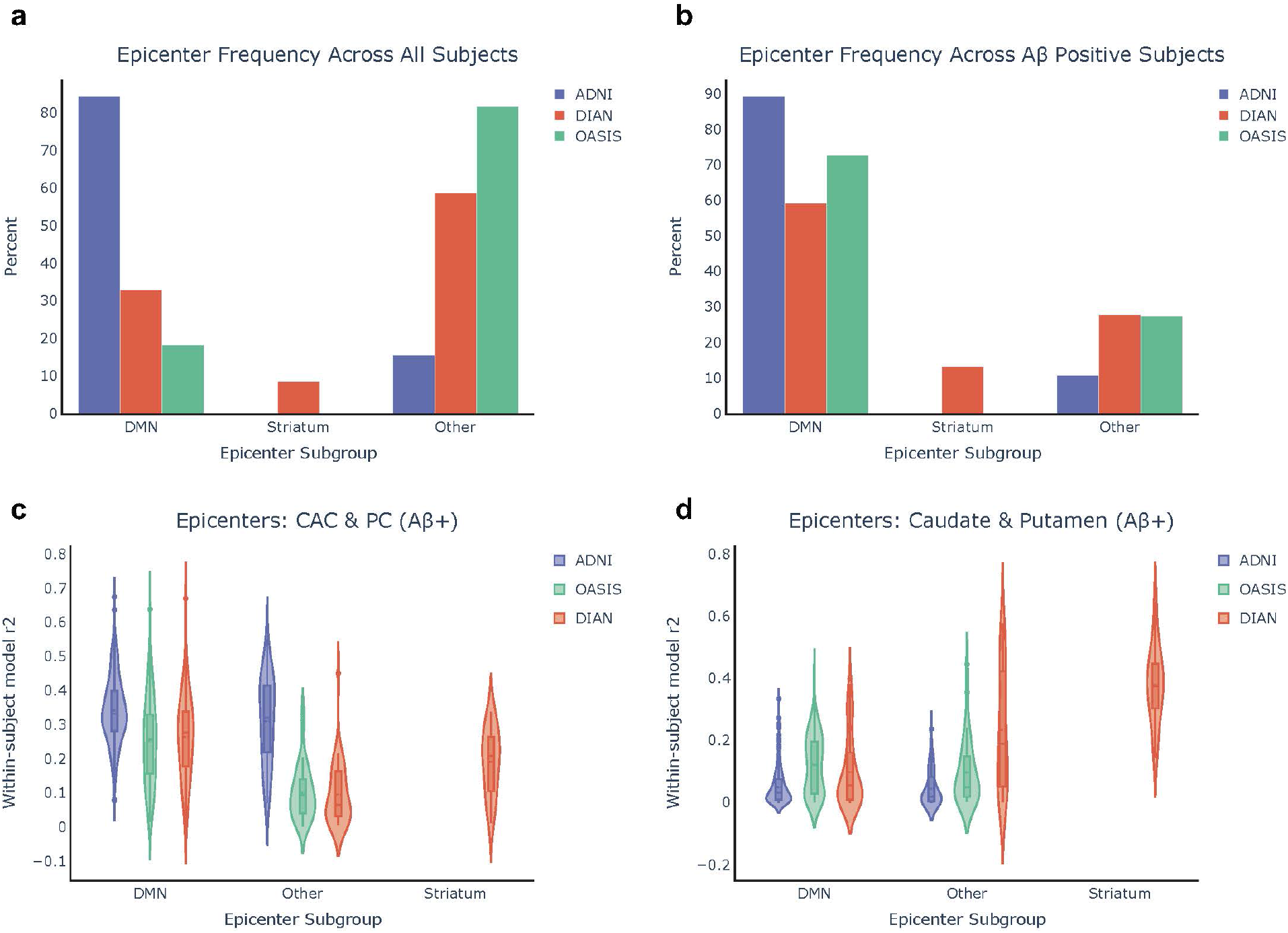
Epicenter frequency and within-subject performance across all datasets. (a) Epicenter frequency across all subjects in each dataset. (b) The same information when only Aβ positive subjects are included from each dataset. (c) and epicenter group, using only Aβ positive subjects. (d) The ESM within-subject performance is shown using the caudate and putamen as epicenters.

We next assessed the performance of the ESM in each “epicenter subgroup” across different model epicenters. We hypothesized that ESM within-subject fit using the caudate and putamen as epicenters would be highest within the DIAN striatum epicenter subgroup, and this was substantiated by the results (Fig 2b). Encouragingly, we found that the ESM within-subject fit using the CAC and PC as epicenters was highest across the DMN epicenter subgroups across all the datasets, and it remained high in the Other subgroup for ADNI. The CAC + PC model fit continued to be higher in ADNI than OASIS (KS=0.42, p=1.6e-12) and DIAN (KS=0.41, p=7.3e-11) within the DMN groups. Within the DIAN dataset, the striatal model significantly out-performed the CAC + PC model in the striatal epicenter subgroup (KS=0.67, p=2.15e-4).

### Epicenter subgroup in DIAN associated with distinct whole-brain Aβ patterns and age at symptom onset

Next, we were interested in parsing the heterogeneity observed within the DIAN dataset with respect to best within-subject epicenter. Specifically, we sought to evaluate any differences in whole brain Aβ pattern and demographics.

As expected, we reaffirmed that individuals in the Other subgroup had significantly lower global cortical Aβ-PET signal (Fig 3c), suggesting these subjects to be ‘false positives’. In other words, individuals with “Other” (i.e. not DMN or striatal) epicenters tended to be low amyloid Aβ-individuals, for whom the model was likely fitting non-specific or off-target binding.

**Figure 3.**
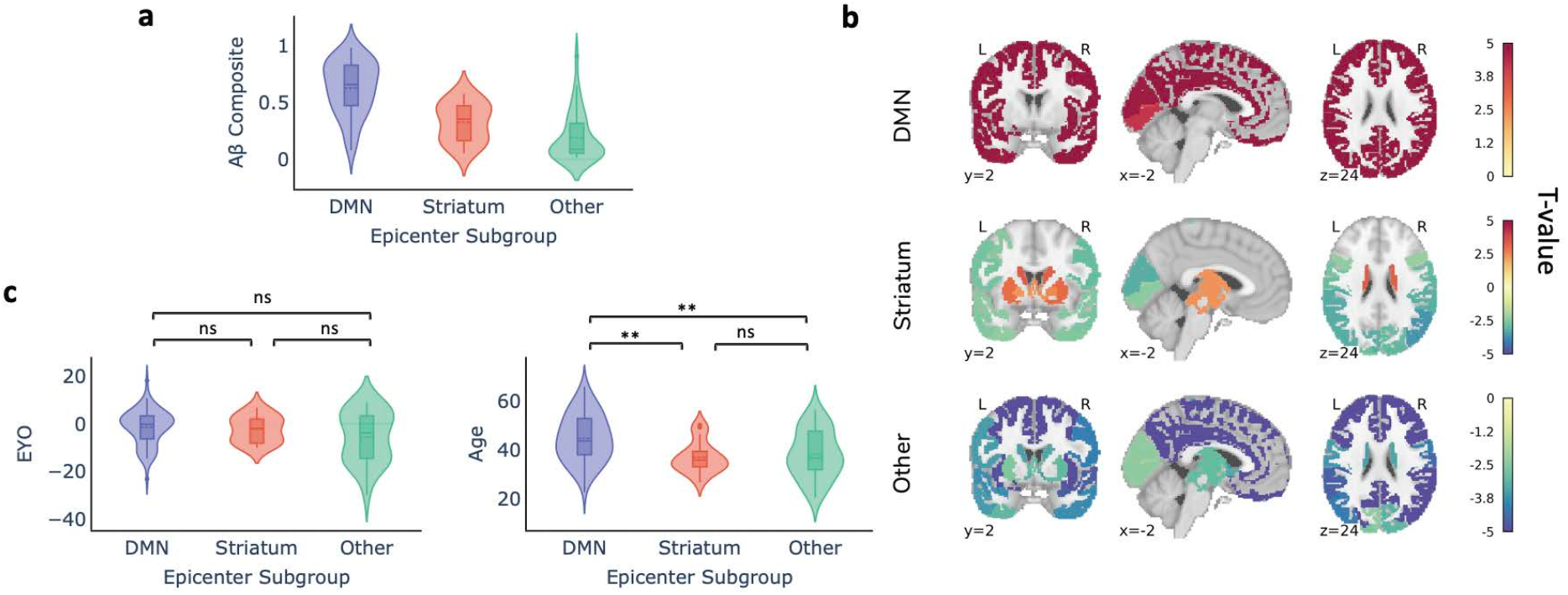
Demographic differences across epicenter subgroups in DIAN (only Aβ positive individuals). (a) Within-subject Aβ composite signal across the epicenter subgroups. (b) Comparison of whole-brain Aβ signal across the epicenter sub-groups. Regions are color-coded based on their t-value for the particular group, with red indicating that there is more Aβ signal in the respective group compared with the other two groups. (c) Within-subject EYO and age differences across the epicenter subgroups.

We further examined whole-brain Aβ pattern differences amongst the different epicenter groups. Individuals whose whole-brain Aβ patterns are best described using a DMN epicenter have more Aβ in the cortex compared with individuals in the other two groups (FDR < 0.05; Figure 3a). Conversely, individuals in the Other epicenter subgroup had less Aβ everywhere in the brain. Individuals with striatal epicenters showed greater striatal PiB binding, but reduced binding in occipital and lateral temporoparietal cortex.

The epicenter groups were also associated with differences in age. Specifically, while the DMN and striatum group did not differ with respect to EYO, individuals in the striatum group were younger than those in the DMN group (Fig 3b). This may potentially suggest that the striatal epicenter phenotype is associated with a younger age at symptom onset and/or an altered disease time course.

### Epicenter reliability across timepoints

With the availability of longitudinal PiB-PET data for a subset of our dataset, we were able to assess how reliably the ESM selects an individual’s epicenter subgroup when presented with data from subsequent timepoints. As shown in Table 1, 124 of the DIAN mutation carriers had two timepoints available, and 44 had three available. Subjects with a DMN or Other epicenter at timepoint 1 (T1) almost always stay that way at timepoint 2 (T2), while there is more variability amongst subjects with a striatal epicenter at T1. This may perhaps indicate that some individuals with a striatal epicenter at T1 are in a temporally short-lived phase whereby Aβ first begins accumulating in the striatum and subsequently in the DMN. In other words, individuals who are advancing with respect to Aβ accumulation may first either show Aβ in the striatum, the striatum then the DMN, or initially in the DMN.

To address this issue of conversion from a striatal epicenter to a different epicenter, we assessed change in composite Aβ deposition probabilities across the different T1-T2 epicenter combinations. We find that individuals who persist with either a striatum or DMN epicenter, or switch from a striatal to DMN epicenter, are gaining amyloid over time (Fig 4b). We observe that those who switch from a DMN or a striatal epicenter to an ‘Other’ are exhibiting a loss of Aβ signal, possibly due to cortical atrophy.

**Figure 4.**
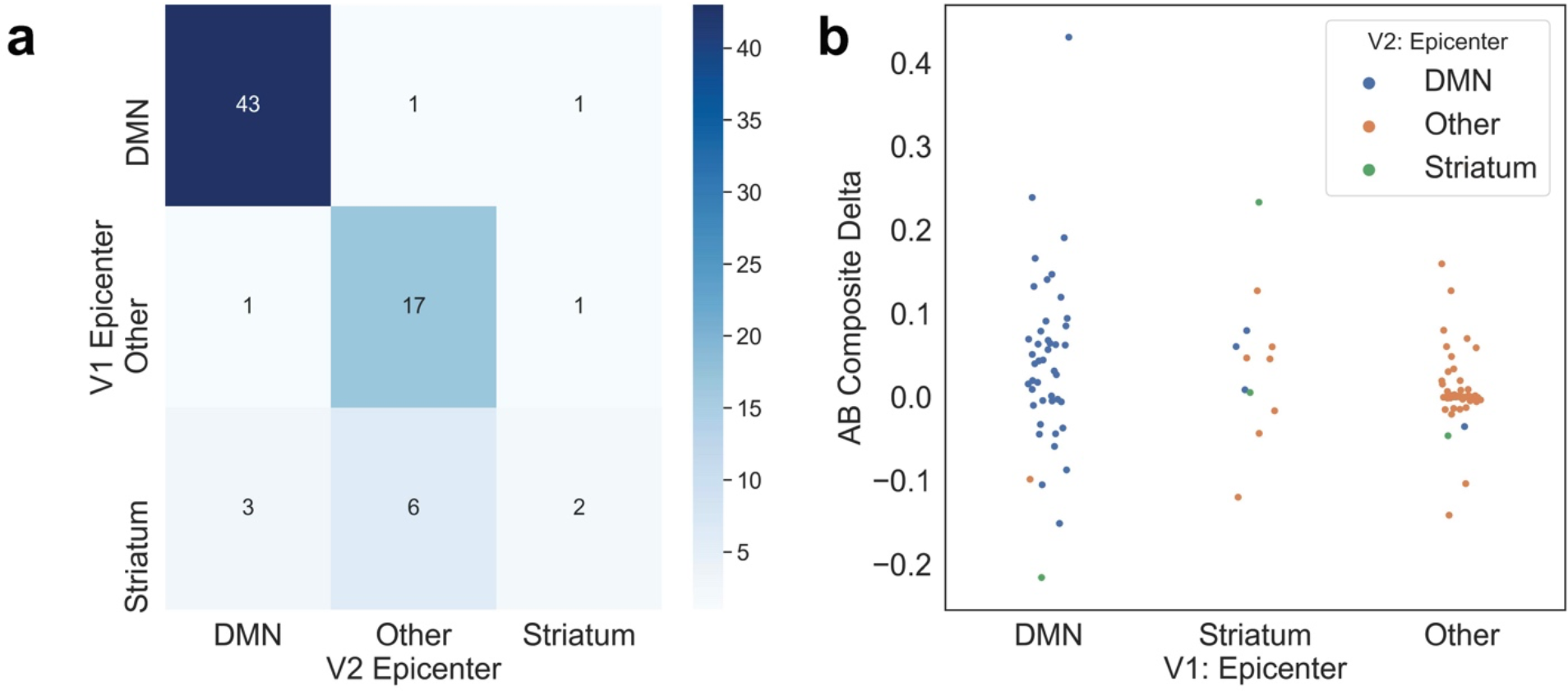
Evaluating epicenter reliability across timepoints in DIAN. A) Confusion matrix for epicenter subgroups at timepoint 1 (T1) vs timepoint 2 (T2). Values along the diagonal represent individuals who remain the same epicenter subgroup at visits 1 and 2. B) Swarmplot representing composite Aβ change in each T1/T2 epicenter subgroup combination.

## Discussion

Throughout this study we have explored how well a model that simulates the transneuronal spread of Aβ under biologically feasible constraints of Aβ production and clearance can explain regional Aβ probabilities for subjects who are either along the sporadic or autosomal dominant Alzheimer’s disease continuum. While many cross-sectional studies have attempted to elucidate differences in the regional Aβ patterns across these subtypes of Alzheimer’s disease, the present study provides a direct comparison of hypothetical spreading patterns of Aβ using a mechanistic model. The ESM generates within subject trajectories of Aβ accumulation, and we leveraged this to assess potential heterogeneity across subjects with respect to the earliest locations of Aβ. Several earlier PiB-PET studies in ADAD have compared which areas begin to accumulate Aβ earliest in the disease time courses of ADAD and LOAD. These studies have reported significantly more amyloid in the striatum in presymptomatic ADAD vs presymptomatic LOAD^5^, and it has been suggested that different mutation types may contribute to heterogeneity amongst individuals with ADAD^7,24^. We found that there was a portion of subjects in the DIAN dataset whose regional Aβ patterns were best reproduced using a striatal epicenter. All but two of these subjects were Aβ positive, suggesting that the results were not driven by false positive signal. Furthermore, these subjects could be distinguished from those with a DMN epicenter by their younger age and younger age of symptom onset, lending support to the idea that this group represents an ADAD-specific phenotype distinct from one characterized by initial Aβ spread from ROIs in the DMN. However, one of the difficulties with interpreting this result lies in the small percentage of subjects with a best fitting striatal epicenter. It is difficult to disentangle whether this striatal epicenter group is truly a separate group for whom Aβ definitively begins accumulating solely in the striatum, or the result of these subjects being imaged during a short dynamic time period, or perhaps both.

However, not all subjects’ Aβ patterns were best recapitulated using a striatal epicenter, and this was supported by the group-level findings. While a hypothetical striatal epicenter explained more variance in the DIAN dataset than in both the ADNI dataset and our validation dataset, OASIS, a DMN epicenter still explained more variance in DIAN within the entire Aβ positive cohort. This may suggest that the Aβ pattern profiles are not homogeneous amongst ADAD mutation carriers, and that there are individuals who are more similar to sporadic AD patients with respect to Aβ. We were able to address this in part by showing that there is a subgroup of DIAN participants whose Aβ patterns are explained as well as the ADNI cohort’s when using the caudal anterior cingulate and posterior cingulate as epicenters.

Our findings provide data-driven corroboration of a neuropathological study finding that ADAD mutation carriers have increased striatal vulnerability to accumulate Aβ due to the regional distribution and metabolism of APP^25^. The same study showed an increased accumulation of striatal tau in ADAD mutation carriers compared with sAD individuals, and previous simulations of tau spreading in sAD shed additional light on how Aβ facilitates the spread of tau and influences its spatial localization^11^. In tandem, a study in ADAD has indicated that striatal amyloid is a better predictor than cortical amyloid of both tauopathy and cognitive decline in ADAD mutation carriers^26^. With availability of tau-PET data for the DIAN cohort, it would be worthwhile to assess this relationship while accounting for the epicenter subgroup differences.

In light of mounting evidence for striatal and network-level involvement in ADAD both with respect to Aβ and tau, a recent study found that frontostriatal circuits are structurally and functionally impacted by APP and PSEN1 mutations^27^. Specifically, the APP gene increased functional connectivity and altered axonal integrity in the caudate to rostral middle frontal gyrus (caudate-rMFG) tract. While the ESM and other mechanistic spreading models reproduce the spread of Aβ over a static network reflecting anatomical connectivity in health, these results, along with those from a separate study evaluating the sequence of changes in anatomical connectivity in elderly individuals’ brains over the course of sAD progression^28^, suggest that Aβ affects the circuits or networks via which it spreads.

One objective of this study was to reproduce the findings in^10^ an independent dataset. One of the issues we observed when modeling group-level results was that of a significant disparity in overall Aβ levels across the three datasets. In particular, the OASIS dataset had a high percentage of younger, cognitively normal adults who were Aβ negative. As we discussed in the Results section, the ESM appears to be sensitive to low levels of Aβ - i.e. the ESM is fit to non-specific or off-target signal not reflecting true pathology, and this would have a particularly large impact on within-subject results for the most likely epicenter(s). As such, we opted to focus on Aβ positive subjects for the within-subject analyses. When we limited our analysis to Aβ subjects, we found that the results across ADNI and OASIS were on par with one another, with a vast majority of subjects being best described by an epicenter that overlaps with the default mode network. This observation is in line with previous datadriven approaches used in both cross-sectional and longitudinal studies to discern which regions begin to show increased Aβ in early stage sAD^29,?^.

This study has several limitations that pertain to measurement of Aβ, anatomical connectivity, and the ESM methodology. One limitation faced when directly comparing the ADNI and DIAN sets is that the PET data was collected using the AV45 radiotracer in ADNI and the PiB tracer in DIAN/OASIS. Additionally, we sought to use the results in the original ESM publication as a benchmark, and this required using the derivatives that had been produced for that paper. Both OASIS and DIAN had been processed using PUP, and there were subsequently differences in the way that the PET scans were corrected for motion and co-registered to the MRI scans. As had been reported in^11^, there are many different choices that can be made in a PET data processing pipeline and the connectivity matrix, and the downstream effects include variable model fit. To determine the best epicenter and by extension epicenter subgroup for each subject, we selected the bilateral ROI that yielded the best within-subject fit, but this method ignores potentially close values across ROIs.

Despite these limitations, our study made several important advances. We show that the majority of Aβ positive subjects in three independent datasets had whole-brain Aβ patterns best reproduced using epicenters overlapping with the DMN. The presence of the younger striatal epicenter subgroup in only the DIAN dataset supports the importance of analyzing differences in individual trajectories, as variability in ADAD disease courses may have important implications for efforts to reduce Aβ burden and improve cognitive impairment.

## Supporting information

Supplemental Figures and Consortium Author List

## Data availability

OASIS-3 and ADNI are open access datasets for which access can be obtained at https://www.oasis-brains.org/ and http://adni.loni.usc.edu/data-samples/access-data/, respectively. The DIAN data can be obtained by request through application, and more information about requesting data access can be found here https://dian.wustl.edu/our-research/for-investigators/dian-observational-study-investigator-resources/data-request-terms-and-instructions/.

## Code availability

The Matlab code for the Epidemic Spreading Model has been made available as a public software release with an accompanying paper (neuropm-lab.com/software^30^). All the Python code used to analyze ESM results, perform statistical analysis, and visualize results can be found at https://github.com/llevitis/DIAN_ESM_AmyloidBeta_Project.git.

## Acknowledgments

We would like to thank Renaud La Joie, Leonardo Iaccorino, Bratislav Misic, and Alain Dager for comments and suggestions during the formulation of this work. Data collection and sharing for the DIAN project was supported by The Dominantly Inherited Alzheimer’s Network (DIAN, U19AG032438) funded by the National Institute on Aging (NIA), the German Center for Neurodegenerative Diseases (DZNE), Raul Carrea Institute for Neurological Research (FLENI), Partial support by the Research and Development Grants for Dementia from Japan Agency for Medical Research and Development, AMED, and the Korea Health Technology RD Project through the Korea Health Industry Development Institute (KHIDI). This manuscript has been reviewed by DIAN Study investigators for scientific content and consistency of data interpretation with previous DIAN Study publications. We acknowledge the altruism of the participants and their families and contributions of the DIAN research and support staff at each of the participating sites for their contributions to this study.

Data were provided [in part] by OASIS OASIS-3: Principal Investigators: T. Benzinger, D. Marcus, J. Morris; NIH P50 AG00561, P30 NS09857781, P01 AG026276, P01 AG003991, R01 AG043434, UL1 TR000448, R01 EB009352. AV-45 doses were provided by Avid Radiopharmaceuticals, a wholly owned subsidiary of Eli Lilly. Data collection and sharing for the ADNI project was funded by the Alzheimer’s Disease Neuroimaging Initiative (ADNI) (National Institutes of Health Grant U01 AG024904) and DOD ADNI (Department of Defense award number W81XWH-12-2-0012). ADNI is funded by the National Institute on Aging, the National Institute of Biomedical Imaging and Bioengineering, and through generous contributions from the following: AbbVie, Alzheimer’s Association; Alzheimer’s Drug Discovery Foundation; Araclon Biotech; BioClinica, Inc.; Biogen; Bristol-Myers Squibb Company; CereSpir, Inc.; Cogstate; Eisai Inc.; Elan Pharmaceuticals, Inc.; Eli Lilly and Company; EuroImmun; F. Hoffmann-La Roche Ltd and its affiliated company Genentech, Inc.; Fujirebio; GE Healthcare; IXICO Ltd.; Janssen Alzheimer Immunotherapy Research Development, LLC.; Johnson Johnson Pharmaceutical Research Development LLC.; Lumosity; Lundbeck; Merck Co., Inc.; Meso Scale Diagnostics, LLC.; NeuroRx Research; Neurotrack Technologies; Novartis Pharmaceuticals Corporation; Pfizer Inc.; Piramal Imaging; Servier; Takeda Pharmaceutical Company; and Transition Therapeutics. The Canadian Institutes of Health Research is providing funds to support ADNI clinical sites in Canada. Private sector contributions are facilitated by the Foundation for the National Institutes of Health (www.fnih.org). The grantee organization is the Northern California Institute for Research and Education, and the study is coordinated by the Alzheimer’s Therapeutic Research Institute at the University of Southern California. ADNI data are disseminated by the Laboratory for NeuroImaging at the University of Southern California.

