## Supplemental Figures and Consortium Author List for "Differentiating amyloid beta spread in autosomal dominant and sporadic Alzheimer’s disease"

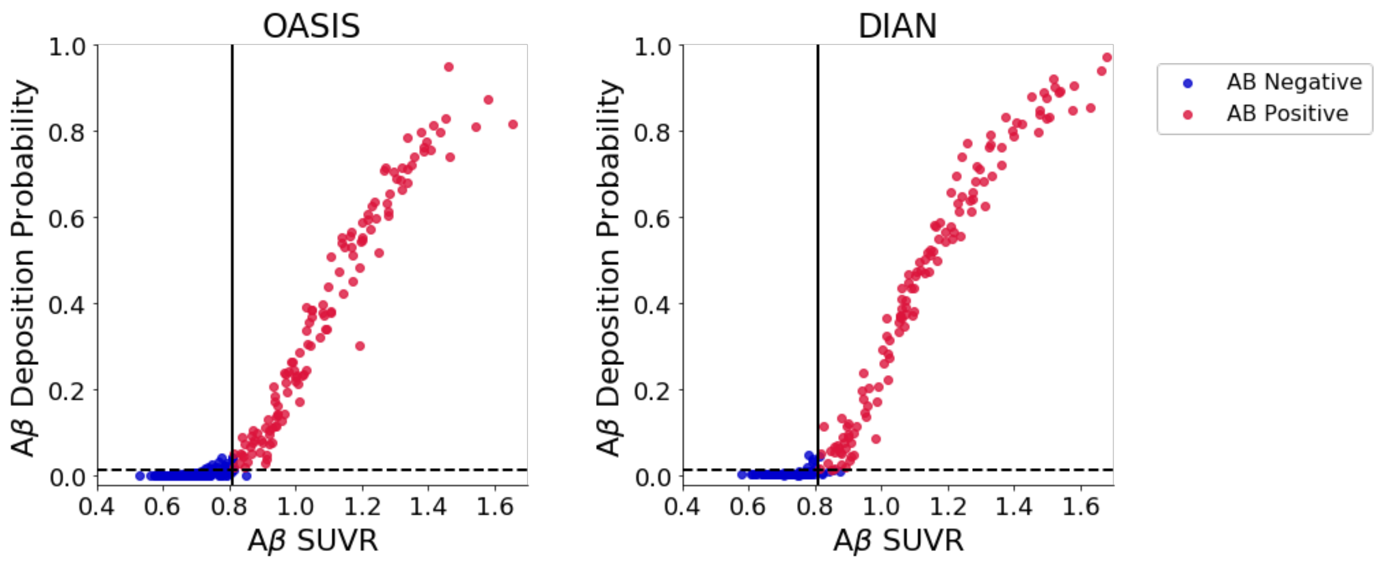


**Supplementary Fig S1**. Aβ positivity defined using joint data driven thresholds computed using SUVR and deposition probability values. The solid line represents the GMM derived Aβ SUVR threshold of 0.81, and the dashed line represents the Aβ deposition probability threshold of 0.0136. Subjects who are positive for both are colored red while those that don't satisfy both criteria are deemed Aβ negative and are colored blue.

### Consortium author list

The DIAN study investigators include: Sarah Adams, MS; Ricardo Allegri, PhD; Aki Araki, ; Nicolas Barthelemy, PhD; Randall Bateman,MD; Jacob Bechara, BS; Tammie Benzinger, MD, PhD; Sarah Berman, MD, PhD; Courtney Bodge, PhD; Susan Brandon, BS; William (Bill) Brooks, MBBS,MPH; Jared Brosch, MD, PhD; Jill Buck, BSN; Virginia Buckles, PhD; Kathleen Carter, PhD; Lisa Cash, BFA; Charlie Chen, BA; Jasmeer Chhatwal, MD,PhD; Patricio Chrem Mendez, MD; Jasmin Chua, BS; Helena Chui, MD; Laura Courtney, BS; Carlos Cruchaga, PhD; Gregory S Day, MD; Chrismary DeLaCruz, BA; Darcy Denner, PhD; Anna Diffenbacher, MS; Aylin Dincer, BS; Tamara Donahue, MS; Jane Douglas, MPh; Duc Duong, BS; Noelia Egido, BS; Bianca Esposito, BS; Anne Fagan, PhD; Marty Farlow, MD; Becca Feldman, BS,BA; Colleen Fitzpatrick, MS; Shaney Flores, BS; Nick Fox, MD; Erin Franklin, MS; Nelly Joseph-Mathurin, PhD; Hisako Fujii, PhD; Samantha Gardener, PhD; Bernardino Ghetti, MD; Alison Goate, PhD; Sarah Goldberg, MS,LPC,NCC; Jill Goldman, MS,MPhil,CGC; Alyssa Gonzalez,

BS; Brian Gordon, PhD; Susanne Gräber-Sultan, PhD; Neill Graff-Radford, MD; Morgan Graham, BA; Julia Gray, MS; Emily Gremminger, BA; Miguel Grilo, MD; Alex Groves, ; Christian Haass, PhD; Lisa Häsler, MSc; Jason Hassenstab, PhD; Cortaiga Hellm, BA; Elizabeth Herries, BA; Laura Hoechst-Swisher, MS; Anna Hofmann, MD; Anna Hofmann, ; David Holtzman, MD; Russ Hornbeck, MSCS, MPM; Yakushev Igor, MD; Ryoko Ihara, MD; Takeshi Ikeuchi, MD; Snezana Ikonomovic, MD; Kenji Ishii, MD; Clifford Jack, MD; Gina Jerome, MS; Erik Johnson, MD, PHD; Mathias Jucker, PhD; Celeste Karch, PhD; Stephan Käser, PHD; Kensaku Kasuga, MD; Sarah Keefe, BS; William (Klunk, MD, PHD; Robert Koeppe, PHD; Deb Koudelis, MHS,RN; Elke Kuder-Buletta, RN; Christoph Laske, PhD; Allan Levey, MD, PHD; Johannes Levin, MD; Yan Li, PHD; Oscar Lopez MD, MD; Jacob Marsh, BA; Ralph Martins, PhD; Neal Scott Mason, PhD; Colin Masters, MD; Kwasi Mawuenyega, PhD; Austin McCullough, PhD Candidate; Eric McDade, DO; Arlene Mejia, MD; Estrella Morenas-Rodriguez, MD, PhD; John Morris, MD; James Mountz, MD; Cath Mummery, PhD;N eelesh Nadkarni, MD, PhD; Akemi Nagamatsu, RN; Katie Neimeyer, MS; Yoshiki Niimi, MD;James Noble, MD; Joanne Norton,MSN, RN, PMHCNS-BC ; Brigitte Nuscher, ; Ulricke Obermüller, ; Antoinette O'Connor, MRCPI; Riddhi Patira , MD; Richard Perrin, MD, PhD; Lingyan Ping, PhD; Oliver Preische, MD; Alan Renton, PhD; John Ringman, MD; Stephen Salloway, MD; Peter Schofield, PhD; Michio Senda, MD, PhD; Nicholas T Seyfried, D.Phil; Kristine Shady, BA, BS; Hiroyuki Shimada, MD, PhD; Wendy Sigurdson, RN; Jennifer Smith, PhD; Lori Smith, PA-C; Beth Snitz, PhD; Hamid Sohrabi, PhD; Sochenda Stephens, BS, CCRP; Kevin Taddei, BS ; Sarah Thompson, PA-C; Jonathan Vöglein, MD; Peter Wang, PhD; Qing Wang, PhD; Elise Weamer, MPH; Chengjie Xiong, PhD; Jinbin Xu, PhD; Xiong Xu, BS, MS;
